# New Histone H4 variant and H2B variant Exhibits Distinct Genomic Locations, Chromatin Affinities, and Dynamics Throughout Life Cycle and Cell Cycle of *Trypanosoma cruzi*

**DOI:** 10.1101/2025.06.20.660671

**Authors:** Juliana Nunes Roson, Mariana Loterio Silva, Herbert Guimarães Silva, Thaina Rodrigues, Nadjania Saraiva de Lira Silva, Camila Gachet-Castro, Héllida Marina Costa-Silva, Vincent Louis Viala, Sergio Shenkman, M. Carolina Elias, Julia P.C. da Cunha

**Author notes:** Address correspondence to Julia Pinheiro Chagas da Cunha.

## Abstract

Histone variants play crucial roles in chromatin organization and transcriptional regulation in eukaryotes. Unusually, trypanosomatids display histone variants for all histones, although a functional homolog of histone H4 variant (H4.V) had not yet been described in *Trypanosoma cruzi*. In this study, we identified a H4.V in *T. cruzi* that is encoded by a single-copy gene located apart from the typical tandem arrays of canonical histone H4. Functional characterization using ChIP-seq assays revealed that H4.V is located at telomeric regions, demarcates convergent strand-switch regions (cSSRs), and determines new transcription termination sites at codirectional PTUs interrupted by tDNA loci. Throughout the cell cycle, H4.V transcript levels remain stable, while protein abundance increases in G2/M, as shown by immunofluorescence and image flow cytometry. In contrast, the histone H2B variant (H2B.V) transcripts peak at S-phase, and protein abundance accumulates progressively. H4.V is more abundant in the nuclei of metacyclic trypomastigotes but barely detectable in amastigotes and bloodstream trypomastigotes. In contrast, H2B.V shows a punctate nuclear pattern and is present in all life stages, with the highest levels also observed in metacyclics. During metacyclogenesis, both variants show a progressive increase in expression, particularly H4.V, suggesting a role in parasite differentiation. Ultimately, salt extraction experiments demonstrated differing chromatin binding affinities across life stages for both variants, with H4.V displaying a more permanent chromatin association than H2B.V, indicating that H4.V may be associated with a more compact chromatin state. Our data reveal H4.V as a novel histone variant in *T. cruzi*, characterized by unique genomic localization, expression profiles, and chromatin-binding dynamics in contrast to H2B.V, underscoring its potential function as an epigenetic marker in transcriptional regulation and adaptation to diverse host environments.

**Author summary:** Chromatin is composed of a group of proteins called histones that are involved in compacting DNA and thereby affecting many DNA-associated mechanisms, such as replication and transcription. Histone variants share distinct primary sequences from the canonical ones and display different roles and genomic locations. While variants of histone H3 and H2A are common in all eukaryotes, variants of H2B and H4 are unusual, except in trypanosomatids, a group of parasites that cause human and veterinary diseases. Among them, *Trypanosoma cruzi*, the causative agent of Chagas disease, exhibits variants for three core histones, but a variant of histone H4 (H4.V) had not been described. Here, we identify H4.V and demonstrate its preferential deposition at telomeres, convergent strand-switch regions (cSSRs), and codirectional polycistronic transcription units (PTUs) interrupted by tDNA loci, likely marking transcription termination sites. Both variants are enriched in the nuclei of metacyclic trypomastigotes; however, H4.V is barely detectable in amastigotes and bloodstream trypomastigotes, while H2B.V is consistently present in all forms. Additionally, each variant shows distinct chromatin-binding affinities across life stages, with H4.V potentially linked to a more compact chromatin state. Our findings reveal a novel histone variant with unique dynamics and epigenetic roles distinct from H2B.V in *T. cruzi*.

## Introduction

Chromatin comprises DNA and proteins that are organized into various structural levels, regulating genome access and, consequently, gene expression (1,2). Histones constitute about 50% of chromatin mass and are classified as either canonical or variant histones. Canonical histones are typically abundant and expressed in the S-phase of the cell cycle; while histone variants differ from their canonical counterparts in their primary sequence, lower abundance levels and the ability to be expressed independently of the S-phase and in a tissue-specific manner (3–5). Canonical and variant histones also differ in their gene and genome organization: while the former are intronless and organized in clusters, the latter are typically encoded by single-copy genes that contain introns (6).

Incorporation of histone variants into chromatin can alter the nucleosome-DNA interactions affecting chromatin organization and accessibility (5–7). Histone variants have been identified across all studied model organisms — from yeast to plants and animals — and for all histone types, although rare examples exist for H2B and H4. Histone variants for histone H3 include the Centromere Protein A (CENPA) - located at centromeres (8) and H3.3 – associated with transcription activation (9). MacroH2A and H2A.X are two histone H2A variants examples, associated with X chromosome inactivation (10) and DNA damage, respectively (11).

Limited H2B and H4 variants were identified in eukaryotes. Among the H2B variants, the H2BT was shown to be associated with spermatid differentiation in mammals (12) and H2B.1 specific to testis, oocytes, and zygotes (13). In the parasite *Plasmodium falciparum* the histone H2B variant (H2B.Z) dimerizes with the H2A variant (H2A.Z) inhabiting the AT-rich promoter regions (14). Regarding histone H4 variants (H4.V), fewer variants were identified, and their functional roles have been largely unexplored. H4 variants were found in *Gallus gallus* (15), in soybeans (16), and recently, a novel Hominidae-specific H4G was described which is predominantly localized in the nucleolus, influencing rRNA expression levels, protein synthesis, and cell cycle progression (17). Examples of H2B.V and H4.V are also described in some trypanosomatids, as discussed below.

Trypanosomatids include some protozoa parasites that cause important medical and veterinary diseases. Among them, the causative agents of Chagas Disease (*Trypanosoma cruzi*), sleeping sickness (*T. brucei*), and leishmaniases (*Leishmania spp.*) are the most studied. These organisms exhibit particularities in their genome organization and gene regulation, such as the presence of polycistronic transcription units (PTUs), intronless genes, and trans-splicing (18). The arrangement of PTUs in the genome gives rise to convergent strand switch regions (cSSRs), where the transcription of two PTUs ends, and divergent strand switch regions (dSSRs), where the transcriptions of two PTUs begins (18–20). These regions encompass, respectively, the majority of transcription terminus site (TTS) and transcription start site (TSS) in trypanosomes.

*Trypanosoma* chromatin is composed of histones that exhibit significant evolutionary divergence, notably in their N-terminal portion (21,22). In contrast to other organisms, histone variants (H2A.Z, H2B.V, H3.V and H4.V) for all canonical histones were found in *T. brucei* (23). Regions associated to transcription initiation and termination are demarcated by histones variants and some histone post-translational modifications (PTMs) across the *T. brucei* genome: H2A.Z, H2B.V, H4K10ac and H3K4me3 are located at TSSs (23–26); while H3.V and H4.V, base J and H3K76me1 and H3K76me2 are enriched in TTSs (23,27,28). H3.V, H4.V and base J are non-essential chromatin factors in *T. brucei*, but knockouts parasites for all three factors exhibit critical changes in cell growth and replication (29). H4.V in *T. brucei* was described to have 86% identity with the canonical H4 (23) and was shown to be the major determinant of transcription termination sites (29).

In the *T. cruzi* genome, all histone variants have been annotated except H4.V; however, only H2B.V has been previously characterized. We have shown that H2B.V localizes at dSSRs and certain tDNA loci. Parasites heterozygous knockout (HzKO) of H2B.V display increased differentiation into metacyclic forms and enhanced invasion of mammalian host cells (30). Intriguingly, H2B.V is overexpressed in tissue-culture-derived trypomastigotes (TCTs) compared to epimastigotes and shows reduced chromatin affinity in TCTs (30). However, it is not known whether *T. cruzi* encodes an H4.V and whether it has a preferential genomic location. Likewise, the expression dynamics and chromatin association of H2B.V and the potential H4 variant across the parasite’s life cycle and cell cycle are still unclear. This study addresses these gaps by identifying and characterizing a novel H4.V in *T. cruzi* that marks cSSRs, some tDNA loci and telomers. We also expand our investigation to better understand the expression patterns and chromatin interactions of both H4.V and H2B.V across life stages and throughout the cell cycle, revealing a variant-specific, dynamic regulatory landscape.

## Results

### In Silico Identification of a Novel T. cruzi H4.V

To search for an H4.V in *T. cruzi* genome, we applied three *in silico* strategies, including analyses of genome locus, alignments, and phylogeny. Histone variant genes are generally dispersed throughout the genome, distinguishing them from the clustered arrangement of canonical histone genes (4,31,32). This dispersed distribution reflects their specialized roles and unique regulatory mechanisms. We confirmed that in *T. cruzi* (CL Brener strain) single-copy of the variant histones H3.V, H2B.V, and H2A.Z are located on chromosomes 40, 27, and 17, respectively, while their canonical counterparts are arranged in tandem on different chromosomes (Fig 1A). For histone H4, we found that most T. cruzi H4 gene copies are located on TcChr2-S (11 copies) and arranged in tandem (Fig 1A), whereas a single-copy gene, TcCLB.511681.20, is located on TcChr24-S, suggesting it may encode a putative H4 variant. The CL Brener strain is a hybrid cell line that displays two haplotypes named Esmeraldo (S) and Non-Esmeraldo (P) like. We found a similar loci distribution of canonical H4 and its putative variant (TcCLB.508739.60) for the non-Esmeraldo like haplotype (8 copies of H4 at TcChr2-P and 1 copy of the putative H4.V at TcChr24-P).

**Fig 1.**
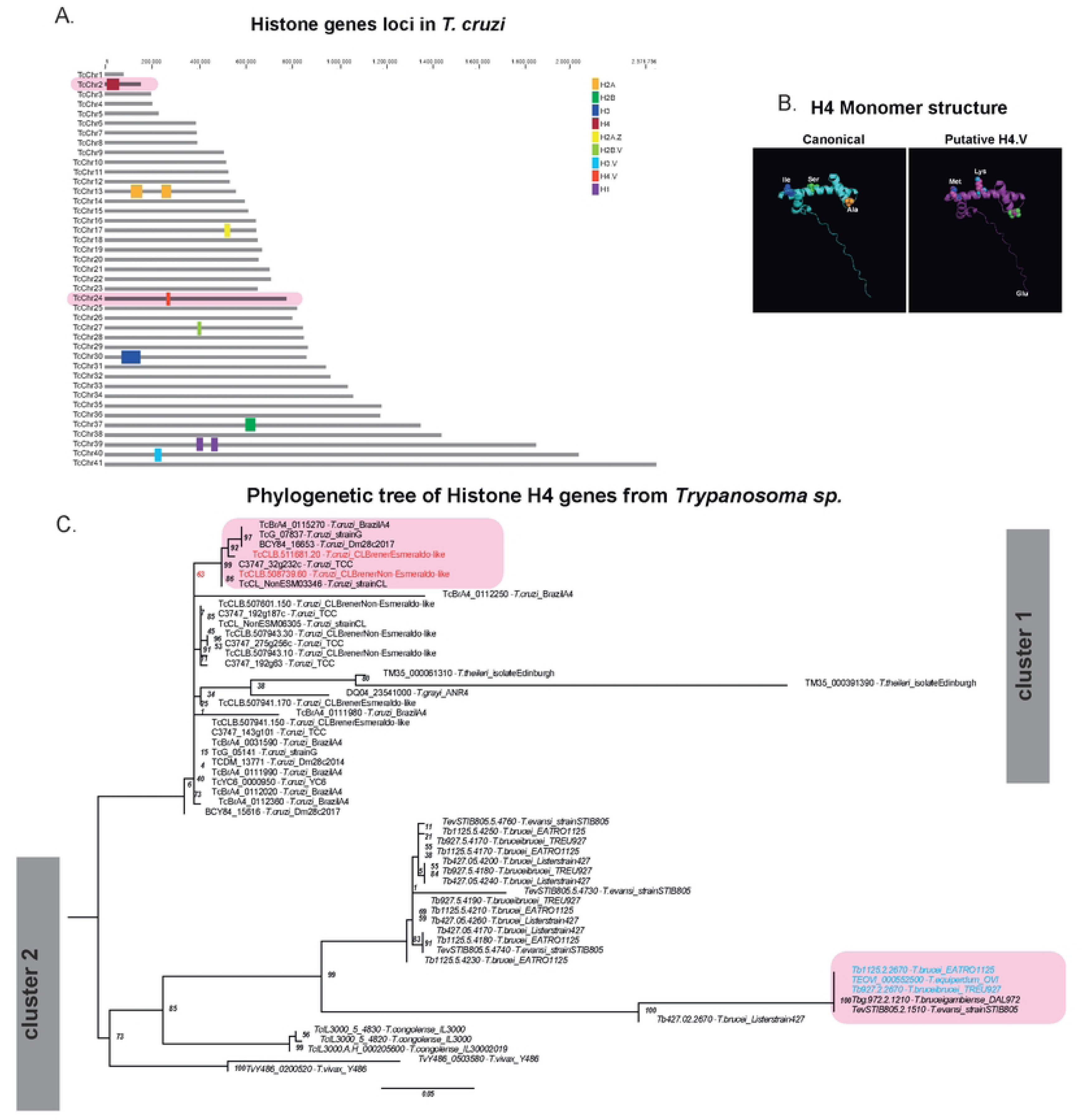
*In silico* identification of putative *T. cruzi* H4.V. **A.** canonical *T. cruzi* (CL Brener strain – S haplotype) histone loci are arranged *in tandem* in specific chromosomes, whereas variants histones are preferentially found in single copies and on different chromosomes. Chromosomes in red highlight canonical histone H4 and the putative H4.V locus. Notably, the histone H1 is the only histone that is not organized in tandem. **B.** Canonical histone H4 and putative H4.V protein structure prediction according *AlphaFold*. Aminoacid differences between two histones are highlighted. **C.** Phylogenetic tree of the H4 histone gene family from the *Trypanosoma* genus. The tree was generated by nucleotide alignments and amino acid 3D structure of histone genes H4 and H4.V obtained from the TriTrypDB database and AlphaFold [80,81]. The species was grouped by the best probability of similarity from the ultrafast bootstrap values (1000 replicates) of the best maximum likelihood (best-fit model (TN+F+G4) - ML) shown at the tree nodes (confidence interval values). The CDSs represented in blue with a pink background are known as histone H4.V. The CDSs in red with a pink background are the putative H4.V from CL Brener strain investigated in this work. For building the tree file, identical H4 sequences of a given strain were considered only once (the identical IDs are listed in S1 Table). The tree layout was edited and exported from the FigTree software program.

Second, by aligning all the H4 sequences (12 sequences from S-haplotype and 9 from P-haplotype (P) (SF1B) we found that canonical histone H4 sequences from both haplotypes are conserved, differing in one amino acid at position 48 (V to I). Interestingly, the putative H4.V single copy-genes shares 98% identity to the remaining histone H4 from CL Brener, differing in three amino acids. The three divergences are: I49M, which preserves the polarity of residues; S59K, which shifts from a polar (serine) to a cationic residue (lysine); and A100E, which shifts from a hydrophobic, nonpolar amino acid (alanine) to one with an anionic charge (glutamine). These amino acid substitutions do not induce major changes on the tertiary structure of the protein (Fig 1B).

Finally, from all histone H4 orthologs of the *Trypanosoma* genus (Fig. 1C, S1 Table), we obtained a phylogenetic tree that evidenced two main clusters composed of histones H4 from *T. cruzi* (cluster 1) and other encompassing sequences from *T. brucei*, *T. congolense,* and *T. equiperdum* (cluster 2), as expected by their phylogenetic differences. Within the second cluster, it is evidenced that known histone H4 variants (highlighted in blue) from *T. equiperdum* and *T. brucei* form a distinct cluster from their canonical counterparts. Within the cluster formed by *T. cruzi* histone H4, it is possible to observe that CL Brener strain sequences are dispersed into many subclusters, although one cluster diverged with 63 node ultrafast bootstrap (confidence interval – in red) from other H4 sequences of *T. cruzi*. Within this cluster, the putative H4.V are found. As a whole, TcCLB.508739.60 (P-haplotype) and TcCLB.511681.20 (S-haplotype) emerge as strong candidates for H4.V in *T. cruzi*.

### *T. cruzi* H4.V demarcates cSSRs, some loci of tDNAs and telomeric regions

To explore the role of this putative histone variant, we used CRISPR-Cas9 to edit its locus obtaining ^Ty1-^H4.V epimastigote lineages (SF2A). The genome editing was confirmed through PCR (SF2B), WB of WCE from epimastigotes (SF2C), and DNA sequencing. We showed that editing the H4.V locus did not affect parasite proliferation or differentiation (SF2D-E).

Considering that histone variants are deposited in specific genomic regions (33), we performed ChIP-seq assays using parasites ^ty1-^H4.V and untagged parasites (Cas9 lineage) as a control. The reads were mapped using genome assemblies available at (34) and at (35) (obtained from long reads), obtaining 74.61% and 85.82% of mapped reads, respectively. We found that the putative H4.V is predominantly enriched in cSSRs (Fig 2A, SF3A-B), as illustrated by the summary plots of all cSSRs (Fig 2B) and polycistronic units (Fig 2C). In the latter, H4.V enrichment is concentrated at the ends of polycistrons, where cSSRs are located. These results indicate that *T. cruzi* H4.V preferentially marks transcription termination regions, similarly to what has been observed for *T. brucei* H4.V [20], suggesting a conserved feature among trypanosomes.

**Fig 2.**
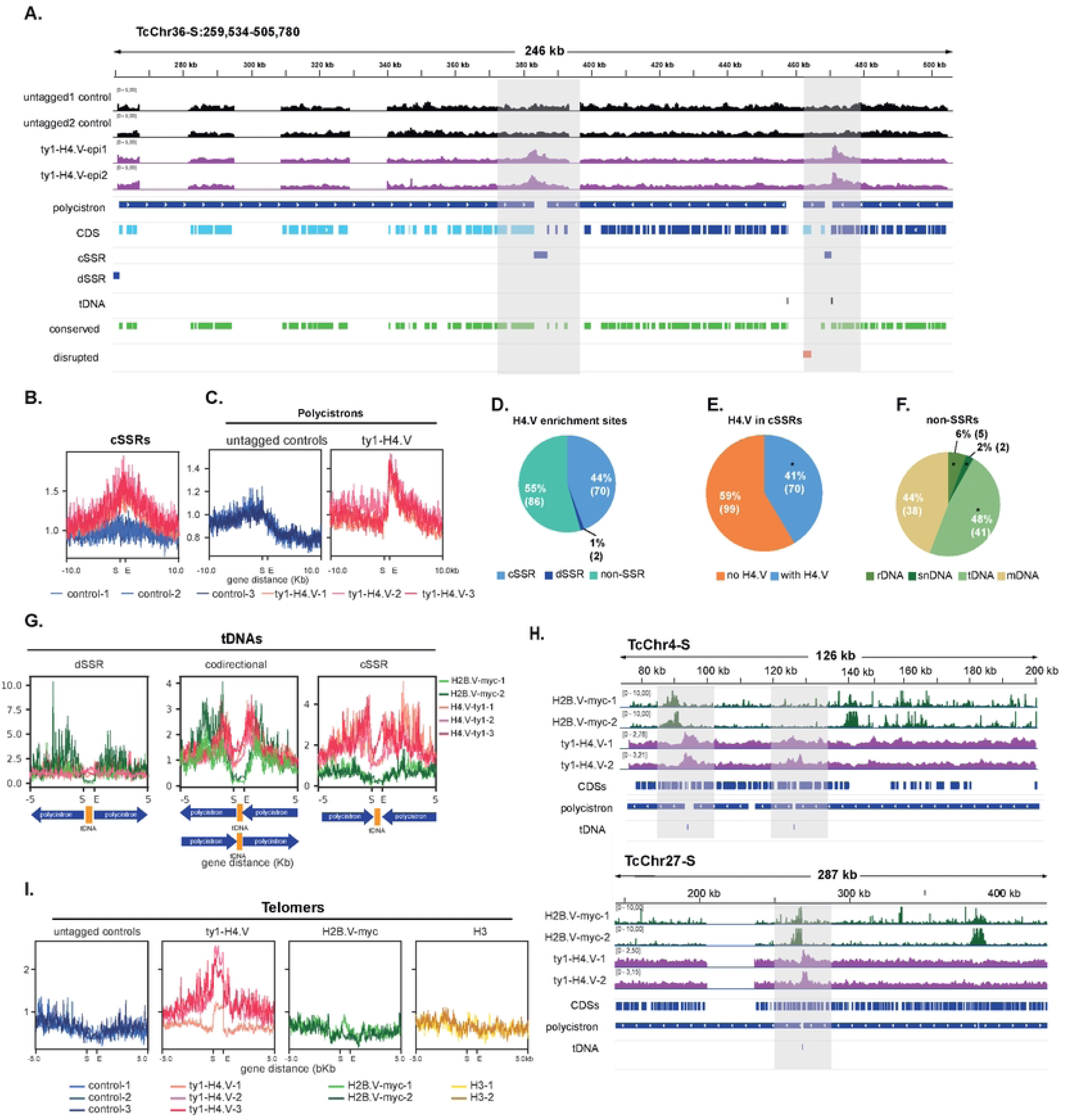
*T. cruzi* H4.V is enriched in cSSRs, near to some tDNAs and telomers. **A.** IGV screenshot of H4.V ChIP-seq data from Chr36. The tracks “^ty1-^H4.V-epi” and “untagged control” represent the read coverage of the ratio values (ChIP/input) obtained by COVERnant (window size of 1001 bases per step of 201). The bed tracks in green represent the conserved genome compartment, in red the disruptive compartment, and in black the tDNA loci. **B-C.** Plot profiles of histone H4.V enrichment detected by ChIP-seq assays in cSSRs (**B.**) and polycistronic units (**C.**). **D.** Pie chart representing the percentage of significant peaks identified by manual curation (Chi-squared test for given probabilities with simulated p-value (based on 1000 replicates) with calculation of the cut-off point for the standardized residues) in genome; in cSSR (**E.**) (cSSR p < 10^15^); in non-SSR (**F.**) (rDNA p<10^15^; snoDNA p<10^8^; tDNA p<10^15^). **G.** Plot enrichment of histones H4.V and H2B.V detected by ChIP-seq assays in tDNAs. The plotted values show that the enrichment of variants does not colocalize in the tDNAs located at dSSRs (8 tDNA genes), but colocalizes at cSSRs (16 tDNA genes), and at codirectional PTUs (31 tDNA genes). **H.** IGV snapshots of two chromosomes harbouring tDNAs located at codirectional PTUs. **I.** Plotprofile of histone H4.V detected in telomeric regions. All summary plots were obtained by the COVERnant ratio values (ChIP/input), and all regions of interest (cSSR, polycistron, tDNA and telomer) were fit into a scale-region represented as “Start” (S) and “End” (E).

Enriched H4.V regions were compiled (SF3C-D and Table S3), revealing that H4.V enrichment is distributed in cSSRs (44%), dSSRs (1%) and non-SSRs (55%) (Fig 2 D). From the 169 cSSRs, 41% of them (n=70) are enriched in H4.V (Fig 2E) (Chi-squared test, p-value <10^15^). When comparing genomic regions adjacent to cSSRs that are either enriched or not enriched in H4.V (SF3A), we observed a notable reduction in the proportion of H4.V in cSSRs between tRNA- and mRNA-coding sequences (from 17% to 0%), and between core regions (from 27% to 13%), suggesting a possible role for H4.V at RNA polymerase II transcription termination sites (TTSs). Conversely, we found an increase in cSSRs between disruptive regions (from 4% to 25%), indicating that these regions may not act as TTSs, or that they rely on an alternative, H4.V-independent mechanism.

Among H4.V enrichment in non-SSRs represented in Fig 2F, 48% represents loci close to tDNAs, 2% snDNAs and 6% rDNAs (Chi-squared test for tDNA p<10^15^, snoDNA p<10^8^, rDNA p<10^15^). Seventy-five percent of tDNAs (41/55) are in regions enriched in H4.V. The frequent localization of H4.V near loci transcribed by RNA polymerases other than RNA Pol II (which drives PTU transcription) suggests the establishment of new TTSs upstream of these regions.

The observation that H2B.V also localizes near tDNA loci (30) raises the question of whether both variants, H2B.V and H4.V, colocalize at the same tDNA. Interestingly, tDNAs located at dSSRs are exclusively enriched with H2B.V, whereas those at cSSRs show exclusive enrichment of H4.V (Fig 2 G). In contrast, tDNAs positioned between codirectional PTUs exhibit enrichment of both variants. Notably, in these regions, H2B.V and H4.V are enriched at distinct positions, possibly marking the new TSSs and TTSs, respectively (Fig 2 G-H).

H4.V is also enriched at the chromosome ends, likely corresponding to telomeric regions (SF4 Fig and S3 Table). Once the telomeric regions are not annotated in available databases, we improved the CL Brener genome assembly using long-read sequencing from (35), resulting on 51 putative telomeric regions (average and median size of 867 and 1024 bp, respectively). We found that telomeric regions are exclusively enriched in H4.V but not in H3 and H2B.V (Fig 2 I). Therefore, H4.V enrichment is detected in all 51 putative telomeres (SF4B).

### Cell Cycle Dynamics of H2B.V and H4.V

*T. cruzi* cell cycle phases display changes in morphology, transcription levels, proteomic and histone PTM patterns (36–39). Using ^[3H]^lysine incorporation into synchronized parasites, it was shown that core histones, along with a subset of histone H1, are synthesized in coordination with DNA replication (S-phase) (40). Histone variants expressions were not evaluated in this study, but they can be expressed and deposited either in a replication-dependent or replication-independent manner (41,42). To investigate the H2B.V and H4.V expression across the cell cycle, we employed a combination of analysis of a public transcriptomic dataset (37), immunofluorescence assays (IFA), and Image Flow Cytometry (IFC).

Cell cycle transcriptomic analysis revealed that H2B.V transcripts peak during S-phase, similarly to the canonical histones H2A and H4, as well as H2A.Z. In contrast, H4.V transcript levels remain relatively stable throughout the cell cycle, with a slight decrease as cycle progresses (Fig 3A). By IFA, H2B.V was consistently detected across all cell cycle phases, with its levels increasing as the cycle progresses (Fig 3B and C). Conversely, H4.V is slightly detect in G1 and S phases but their levels were more prominent in G2 and G2/M (Fig 3D and E). H4.V increased levels in later cell cycle phases were confirmed by IFC assays (Figs 3E and S5). Together, results revealed a progressive increase in both histone variants as cycle progresses.

**Fig 3.**
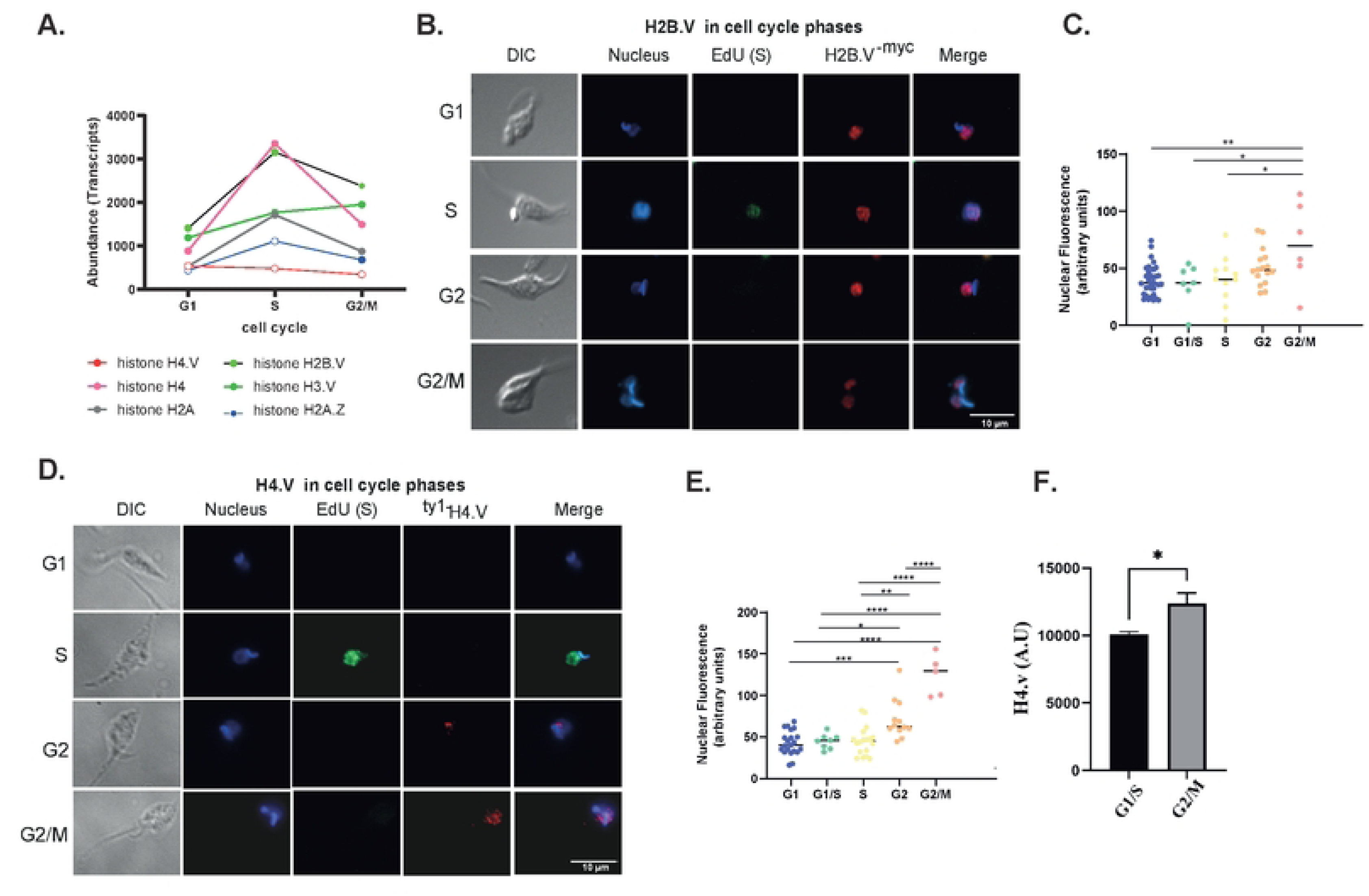
Location and abundance of histone variants in the *T. cruzi* cell cycle. **A.** Quantification of histone transcripts abundance from (37) cell cycle transcriptomic analysis of *T. cruzi*. IFA of H2B.V^- myc^ (**B.)** and ^ty1-^H4.V (**D.)** parasites from EdU incorporation assays in G1, S and G2/M cell cycle phases. Quantification of nuclear labelling from IFA of H2B.V^-myc^ **(C.)** and ^ty1-^H4.V **(E.)** parasites from two biological replicates. For H2B.V^-myc^ and ^ty1-^H4.V lines, 136 and 102 parasites were evaluated. Quantifications were performed in ImageJ considering equal nucleus area. Ordinary one-way ANOVA with Tukey’s multiple comparisons test, where * (p<0.05); and ** (p<0.005). **F.** Histogram representing the median intensity values of ^ty1-^H4.V labelling in G1/S and G2/M from IFC data. Cell cycle determination was based on nuclear/kinetoplast areas obtained by DNA labelling (DRAQ5 detection) (SF5). Upon filtering events, the estimated cell cycle gates were obtained (SF5 B) (see gating strategies at M&M section).

### H2B.V and H4.V exhibit Specific Expression Patterns Across *T. cruzi* Life Stages

*T. cruzi* undergoes morphological and gene expression changes between replicative (epimastigotes and amastigotes) and non-replicative (trypomastigotes – TCT and metacyclics) stages, adapting to distinct microenvironments (43–47). Life form transition is also followed by alterations in nucleus and chromatin structure, including histone PTMs (48–51). Given that the histone variants H2B.V and H4.V exhibit distinct characteristics mainly related to genomic localization and cell cycle expression, we questioned whether their distribution and abundance might also vary within the nucleus during life cycle stages.

H2B.V labelling was observed in the nuclei of all *T. cruzi* life forms by IFA (Fig 4A). This variant histone displays a punctate pattern distributed throughout the nucleus, excluding the nucleolar region. In the infective forms (TCT and metacyclic forms), the H2B.V staining resembles that in epimastigote forms but appears more dispersed, likely due to the elongated nuclear morphology of these forms. While H4.V showed more intense labelling in the metacyclic nuclei compared to other life forms. In epimastigotes, H4.V labelling varies in accordance with the cell cycle, as we have seen above. In contrast, H4.V is minimally detected in amastigotes and TCT forms in IFA assays. Unlike the punctate pattern observed for H2B.V, H4.V appears uniformly dispersed throughout the nucleus, with some metacyclic forms displaying a stronger signal.

**Fig 4.**
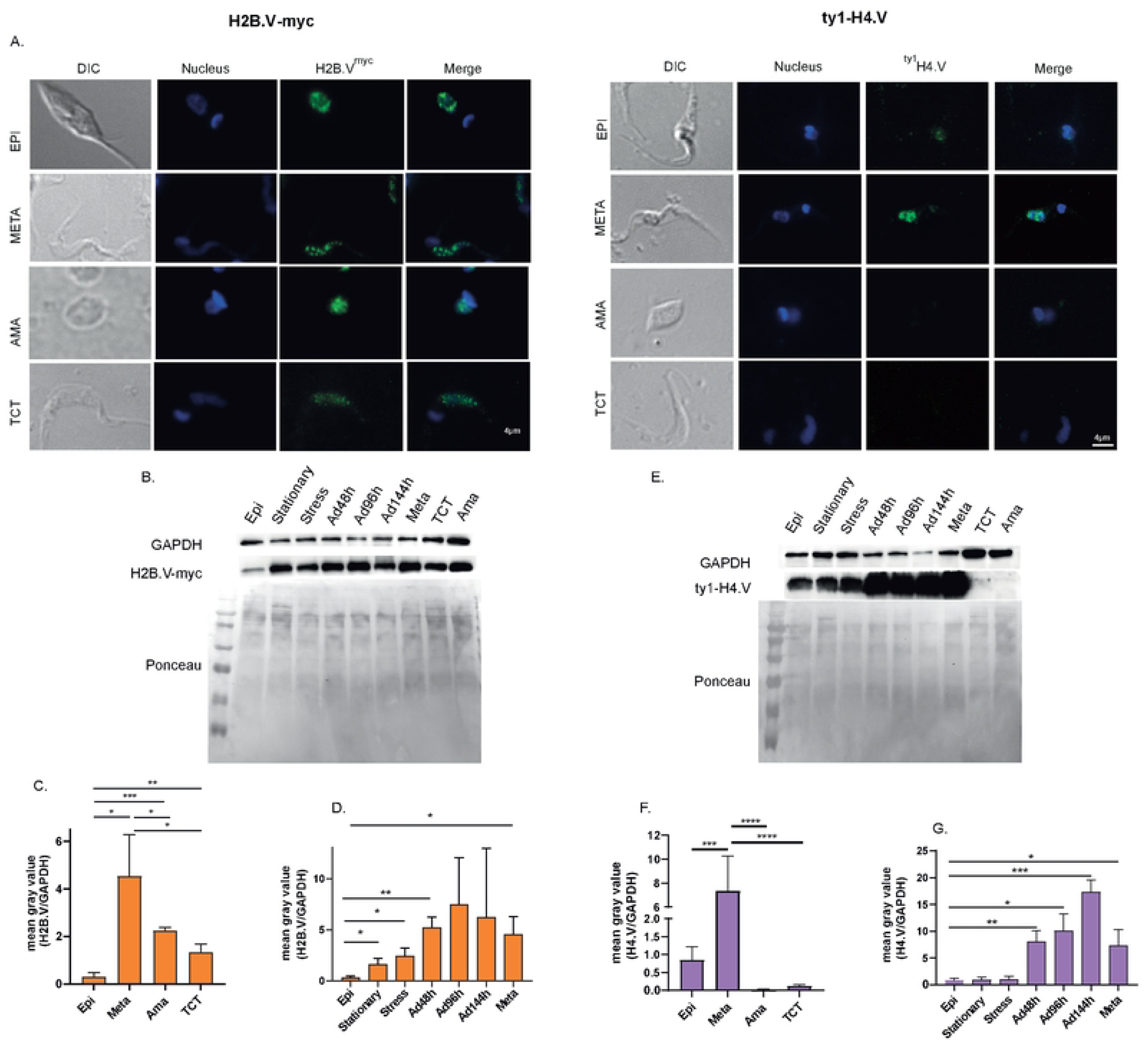
Subcellular localization and abundance of histone variants among *T. cruzi* life forms. **A.** Cellular localization of H2B.V and H4.V on different life stages of *T. cruzi*. Immunofluorescence of the parasites expressing either H2B.V^-myc^ (left) or ^ty1-^H4.V (right) in different life stages of *T. cruzi*. **B.** WB assays of WCE of parasites expressing H2B.V^-myc^ during metacyclogenesis and life forms. **C-D.** Quantification (mean grey value) of H2B.V abundance normalized by GAPDH abundance obtained from (B.) showing significant differences of H2B.V among life forms (C) and metacyclogenesis (D). **E.** WB assays of WCE of parasites expressing ^ty1-^H4.V during metacyclogenesis and life forms. **F-G.** Quantification (mean grey value) of H4.V abundance normalized by GAPDH abundance obtained from (E.) showing significant differences of H4.V among life forms (F) and metacyclogenesis (G). Each well contains WCE of 5x10^6^ parasites. For all quantifications, we used an unpaired t-test of multiple comparisons, in which * = p<0.05; ** = p<0.01; *** = p<0.005; and **** = p<0.001. EPI = epimastigote; META = trypomastigote metacyclic; AMA = ama-like (extracellular amastigote); and TCT= tissue-culture-derived trypomastigote.

By WB we detected that H2B.V abundance is more pronounced in TCT compared to epimastigote forms (30). Here, we expand this analysis comparing the H2B.V abundance among all life forms (Fig 4B and C). Hence, H2B.V is more abundant in metacyclics, followed by amastigotes, TCTs and epimastigotes. Furthermore, changes in H2B.V abundance were detected along metacyclogenesis, mainly at initial stages of differentiation (epimastigote vs. stationary epimastigote vs. stress vs. Ad48h-p-value < 0.05) (Fig 4 B and D).

In line with the predominant H4.V labelling in metacyclic forms observed by IFA (Fig 4A), WB assays confirmed a significantly higher H4.V abundance in these forms (Figs 4E and F). H4.V abundance shows a progressive increase throughout metacyclogenesis, as demonstrated in Fig 4G (unpaired multiple t-test, Epi stationary phase vs. ad144h, *P* < 0.0005). The presence of H4.V in epimastigotes, its increased expression in metacyclics, and its low expression levels in amastigotes and TCTs suggest that H4.V expression may be associated with the microenvironment of the invertebrate host likely affecting expression of surface virulence factors.

### Histone variants Chromatin Binding Affinities Across *T. cruzi* Life Stages

Histone variants and canonical histones can differently impact chromatin structure, either promoting a more accessible or stable chromatin state (52). Previously, we observed that H2B.V and H3 are released from chromatin at lower salt concentrations in TCTs compared to epimastigote forms, suggesting that these histones have a weaker binding affinity to TCTs’ chromatin (30). In this study, we extend this analysis by investigating whether histone variants (H2B.V and H4.V) and a canonical histone (H3) display differential chromatin binding affinities across distinct life stages.

Fig 5A demonstrates that H2B.V is consistently released into the cytoplasmic fractions (S1 and S2 Figs) across all three life forms analysed, while H4.V is released only in the epimastigote forms (Fig 5B). Histone H3 is released in the cytoplasmic fractions of epimastigotes and TCTs. These findings suggest that each histone has an unbound chromatin fraction that can vary among life stages. When comparing chromatin affinity across life forms, we observed that in epimastigotes and metacyclics, H2B.V begins dissociating from chromatin at 400 mM NaCl, retaining most of its interaction with chromatin. The canonical histone H3 follows a similar pattern, starting to dissociate at 400 mM NaCl, with a significant fraction remaining chromatin associated. In contrast, H4.V remains bound at chromatin even when submitted to 600 mM NaCl. These data suggest that in epimastigotes and metacyclics, H2B.V has the weakest chromatin interaction, followed by H3, with H4.V showing the strongest binding.

**Fig 5.**
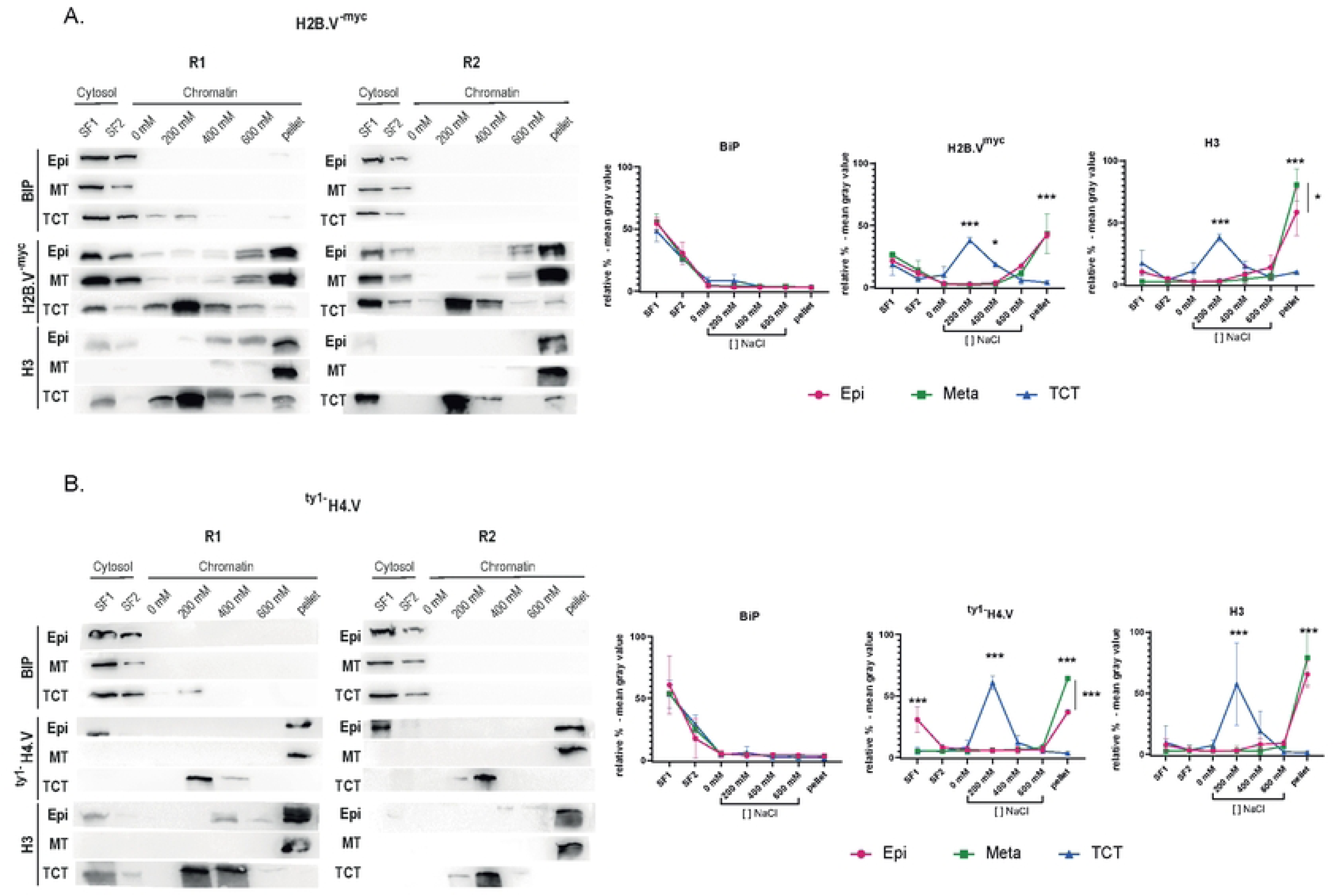
Assessment of the relative chromatin affinity of H2B.V, H4.V and canonical histone H3 in epimastigotes, metacyclics and TCTs life forms. Cytosolic (SF1 and SF2) and chromatin-soluble fractions from saline extraction (0, 200, 400 and 600 mM) of H2B.V^-myc^ (A.) and ^ty1-^H4.V (B.) parasites were obtained and evaluated by WB. WB were performed in biological replicates using antibodies against BiP (cytosol marker), myc (H2B.V), ty1 (H4.V) and histone H3. Protein amounts relative to equal amounts of DNA were applied to each well, fractionated in 15% SDS-PAGE. On the right, quantification (mean gray value) of the WB assays. ANOVA Tukey’s multiple comparisons test among life forms: * p<0.05 and *** p<0.0001.

In contrast, TCT’s chromatin exhibits markedly lower histone interaction affinity. All histones analysed dissociate at 200 mM NaCl, with a near-complete release occurring at 400 mM NaCl. This indicates that TCT’s chromatin is structurally distinct from that of the other compared life forms, particularly in terms of histone-binding affinity.

## Discussion

In this study, we identified a novel histone H4 variant and characterized its genomic localization, abundance, and chromatin affinity across the *T. cruzi* life cycle and cell cycle. Evaluating the also variant H2B.V, we found that the two histone variants share both similar and different features related to their expression/abundance, genome location, and chromatin-binding affinities in *T. cruzi* life and cell cycle. The H4.V identification was supported by four main lines of evidence: (i) divergences in sequence and phylogeny compared to canonical H4; (ii) localization at unique genomic loci, distinct from the tandemly repeated canonical H4 sequences; (iii) stage- and cell cycle-specific abundance, suggesting specialized regulation; and (iv) specific genomic location at cSSRs, near to some loci of tDNAs – likely demarcating TSSs, and at telomeric regions. Notably, H4.V localization at cSSRs contrasts with the preferential association of H2B.V with dSSRs (23). These conserved genomic location distributions in trypanosomes highlight a potentially conserved role for histone variants in transcription initiation and termination regions in these organisms.

It is known that H2B.V–H2A.Z dimers are more unstable than canonical H2B–H2A dimers (24), likely contributing to a more open chromatin configuration that is permissive to transcription initiation. Here, we showed that H2B.V exhibits weaker chromatin interaction compared to H3 and H4.V, supporting the presence of a more unstable chromatin structure at TSSs, in contrast to a more compact and stable chromatin at TTSs. The latter likely forms a structural barrier that halts RNA polymerase II progression into adjacent regions, thereby preventing transcriptional readthrough. In eukaryotes, chromatin structure plays a role in slowing down transcription termination (53). In trypanosomes, TTSs are enriched in base J, a modified thymine that is important for RNA polymerase II termination, as its absence leads to transcriptional readthrough (54). Studies in *T. brucei* using parasites knockouts for all three TTS marks (H3.V, H4.V and base J) identified H4.V as a major signal for transcription termination (29). The predominant localization of H4.V at cSSRs, which are likely to colocalize with base J and H3.V in *T. cruzi*, suggests that these epigenetic marks may act together to promote transcription termination. Since all marks are deposited following DNA replication, the question remains as to which is deposited first and whether one influences the recruitment of the other — a topic that warrants further investigation.

Aside from H4.V enrichment at cSSRs, we also found depositions of both H4.V and H2B.V in PTUs that are interrupted by loci of tRNAs, implying the creation of new TTSs and TSSs into non-SSRs. The H4.V enrichment in telomeric regions is particularly noteworthy, as *T. cruzi* telomeric and subtelomeric regions are known to house multigene families (55) associated with surface virulence factors (trans-sialidases, mucins, MASP, Gp63) and chromatin structure regulation (RHS) (56). This pattern, together with the potential colocalization of H4.V and base J (23,57) in these regions, seems to be specific to *T. cruzi*, since in *T. brucei*, H4.V is absent from telomeric regions, where H3.V is instead enriched (23). Notably, H3.V deletion in *T. brucei* disrupts VSG silencing (29), suggesting a role for histone variants in the regulation of sub-telomeric gene expression. Altogether, these findings support the idea that histone variants H4.V and H3.V may have regulatory roles in the expression of virulence genes located in sub- and telomeric regions. Future studies using *T. cruzi* H4.V knockout lines will be essential to determine whether this variant directly contributes to the regulation of virulence factor gene expression.

H4.V and H2B.V exhibit distinct abundance patterns throughout the parasite’s life cycle. Both variants are present in epimastigotes and metacyclic forms; however, H4.V is nearly absent in intracellular amastigotes and bloodstream trypomastigotes. This observation suggests a specialized role for H4.V in the insect vector environment or that H4.V expression is essential for metacyclogenesis likely affecting surface virulence expression. During metacyclogenesis, H4.V levels progressively increase, whereas H2B.V undergoes a pronounced upregulation early in the differentiation process. Notably, the levels of both variants are higher in metacyclics compared to epimastigotes. Metacyclic forms are characterized by reduced transcriptional activity and more condensed chromatin dispersed throughout the parasite nucleus (50,51). The enrichment of H4.V in telomeres, known to be enriched in heterochromatin (58) along with its higher chromatin affinity, suggests that this variant is associated with a more compacted chromatin state. Whether H4.V is directly involved with the generation of a more compacted chromatin state needs to be further evaluated, as well as the impact of H4.V higher enrichment in metacyclics. Further studies, such as ChIP-seq assays, are needed to confirm whether the genomic localization observed in epimastigotes is preserved in metacyclics or if these variants are redistributed more broadly across the genome, potentially leading to the increased global levels of H4.V in these forms.

On the other hand, the enrichment of H2B.V in metacyclics appears counterintuitive, as nucleosomes containing H2B.V-H2A.Z dimers are typically associated with nucleosome destabilization and enhanced chromatin accessibility (23). Consistent with this, we observed that H2B.V interacts weakly with chromatin in both epimastigotes and metacyclics. Therefore, we hypothesize that the increased abundance of H2B.V in metacyclics may reflect chromatin remodelling processes rather than direct association with transcriptional activation.

Finally, our findings suggest potential implications for histone variants in the three-dimensional organization of chromatin. Recently, we found that cSSRs and tDNA loci of *T. cruzi* are preferentially located at the borders of chromatin-folding domains and evidenced that some tDNA loci interact with each other in a 3D organization (59). Knockouts for H4.V and H3.V in *T. brucei* displayed critical changes in 3D organization (29). Altogether, the deposition of T. cruzi histone variants at key genomic regions—such as dSSRs, cSSRs, tDNA loci, and telomeres—which occupy strategic positions in the 3D genome, suggests that these variants contribute to the establishment and/or maintenance of global chromatin architecture. Additionally, their varying abundance and chromatin-binding dynamics throughout the life cycle, along with their enrichment at sites of transcription initiation and termination, suggest that transcriptional may also be modulated by the dynamic behavior of these histone variants.

Overall, our study emphasizes the complex and dynamic roles of histone variants in *T. cruzi*, opening avenues for future research into their functional contributions to chromatin structure and transcriptional across the parasite’s life cycle. Comparison of H4.V and H2B.V features underscored the conserved role of histone variants in critical transcription initiation and termination regions in trypanosomes as well as highlighting their specific features in cell cycle and life forms.

## Materials and Methods

### Phylogenetic Analysis – Cladogram of Histone H4 Genes in *Trypanosoma spp*

*Trypanosoma spp.* H4 coding sequences (CDS) from the TriTrypDB database (https://tritrypdb.org/tritrypdb/app - downloaded at September 2022) were compiled into a FASTA file. Identical H4 sequences per strain were considered only once (S1 Table). Sequences exceeding 300 bp were eliminated from the dataset, realigned by MAFFT v7.475 program (60), and adjusted by manual curation with Aliview (61). Translated sequences were aligned with predicted structural sequences from AlphaFold (“Q4CMF1.pdb” – putative H4.V and “Q4DDL6.pdb” – canonical H4) using PROMALSD3 (62). Subsequently, the structural sequences obtained were excluded from the 3D alignment file and realigned with CDS nucleotides using the RevTrans 2.0 tool (63) to generate the final cladogram dataset. The maximum likelihood (ML) phylogenetic tree was estimated using the IQ-TREE program (64), applying the ultrafast bootstrap statistics algorithm with 1000 replicates to find the best-fit model. The found best-fit model was applied in a second phylogenetic tree performance. Tree files were analysed using the software FigTree v1.4.4 (http://tree.bio.ed.ac.uk/software/figtree/).

### Parasite Cultures

*T. cruzi* CL Brener epimastigote forms were cultured in complete LIT medium (65) supplemented with 10% fetal bovine serum (FBS) and 0.5% hemin at 28°C. The constitutive Cas9 lineage (66), was supplemented with G418 (100 μg/mL); the ^ty1-^H4.V and H2B.V^-myc^ lineages with blasticidin (10 μg/mL) and puromycin (5 μg/mL), respectively. Metacyclic trypomastigotes were obtained following the protocol described by (67) with some alterations. The epimastigotes in stationary phase were incubated for 2 h in TAU medium(190 mM NaCl, 17 mM KCl, 2 mM CaCl_2_, 2 mM MgCl_2,_ 8 mM sodium phosphate buffer, pH 6.0) at 5 x 10^7^ parasites/mL, followed by dilution in TAU3AAG (TAU medium supplemented with 20 mM L-proline, 10 mM glucose, 50 mM glutamic acid, 2 mM aspartic acid) with 5 x 10^7^ parasites/mL. The cultures were then incubated horizontally at 28°C with 5% CO_2_ for 144 hours. Metacyclic purification was performed following the protocol described by (68). *T. cruzi* trypomastigote forms (TCTs) were obtained from the supernatant of infected mammalian cells (LLCMK2) maintained in DMEM medium supplemented with 10% FBS at 37°C and 5% CO_2_ as described by (69). After the 15th day of infection, extracellular amastigote forms were obtained and the released TCTs were incubated in the complete LIT medium for 24 hours to differentiate into amastigote-like forms (70). Cells were counted using Neubauer chambers in biological triplicates.

### Growth Curve

A total of 5 × 10^6^ epimastigotes/mL were transferred to a 24-well plate, and their growth was monitored over a period of 4 days. Parasite counts were conducted using a Neubauer chamber, and the means and standard deviations of the replicates were analysed relative to the growth of Cas9 lineage parasite (control).

### ^ty1-^H4.V parasite generation

^ty1-^H4.V lineage generation was obtained by CRISPR/Cas9 as described before (30,71) with some alterations. DNA from the plasmid pPOTv6-blast-mNG, containing the blasticidin resistance gene and the ty1 tag gene, along with the sgRNA targeting the 5’ UTR region of H4.V were amplified using Phusion™ High-Fidelity DNA Polymerase (ThermoFisher). The primers “Donor-H4V-Fw” and “Donor-H4V-Rv” were utilized to amplify the donor DNA from the plasmid pPOTv6 -blast-mNG. The PCR product comprised 30 base pairs of the homologous arm (the initial 30 nucleotides of the H4.V gene - (TcCLB.511681.20), excluding the methionine codon), three copies of the Ty1 tag sequence, the blasticidin resistance gene, and another 30 bp of homology arm (30 nucleotides from the intergenic region adjacent to the cleavage region). The donor DNA was inserted into the 5’ region of the H4.V locus for expression in the N-terminal portion of the H4.V protein. Primers “sgRNA-H4V-146” and “sg scaffold” were employed to amplify the sgRNA. PCR products were purified using the QIAquick PCR Purification Kit and transfected into epimastigotes expressing Cas9 (CL Brener strain). Transfections were carried out using the same buffers described for the generation of H2B.V^-myc^ parasites (30), employing the Amaxa Nucleofactor electroporator (Lonza) with program X-014 in 0.2 cm cuvettes. The transfected parasites were then transferred to 5 mL of complete LIT medium and incubated at 28°C. After 24 hours, the cultures were supplemented with 10 μg/mL of blasticidin, and selected transfectants were maintained for two weeks. Parasite clones were obtained by limiting dilution with conditioned medium with 10 μg/mL blasticidin. Genome editing was confirmed by PCR using primers Fw-UP-blast-check (1), Fw-UP-^ty1-^check (2), H4V_Int3_Rv-check (3), and H4V_Rv-check (4), with primer sequences detailed in S2 Table.

### ChIP-seq ^ty1-^H4V

ChIP-seq experiments were performed following the protocol outlined in (30) with certain modifications. We used 3x10^8^ of epimastigote forms (^ty1-^H4.V and untagged Cas9 lineage parasites) in biological triplicates. For the sonication, the samples were resuspended in TELT buffer (50 mM Tris–HCl pH 8.0, 62.5 mM EDTA, 2.5 M LiCl, 4% Triton X-100) and sonicated in the Q800R3 sonicator (Qsonica) under the following conditions: 50% amplitude, 15 seconds on, 30 seconds off for 30 minutes at 4°C. The Dynabeads protein G (Thermo Fisher) ressuspended in blocking buffer (5 mg/mL of BSA diluted in PBS) containing 2.3 µg of anti-ty-1 antibody (MA5-23513 - Invitrogen), was added to the supernatant and incubated at 4°C overnight with slow rotation. Following incubation, washes were performed with ice-cold RIPA buffer (50 mM HEPES pH 7.6, 1 mM EDTA, 0.7% sodium deoxycholate, 1% NP-40, 0.5 M LiCl, and 1 mM PMSF), followed by a final wash with TE buffer (10 mM Tris-HCl pH 8.0 and 1 mM EDTA pH 8.0). The chromatin was eluted in elution buffer (50 mM Tris-HCl pH 8.0, 10 mM EDTA pH 8.0, and 1% SDS) at 65°C for 30 minutes. After DNA purification, described in (30), ChIP and input samples were used to construct Illumina DNA Prep (M) Tagmentation libraries using indexes adapters (IDT for Illumina DNA/RNA) and subsequently sequenced using Illumina NextSeq 1000/2000 (100 bp - pair end).

### ChIP-seq Data Analysis

ChIP-seq processing and analysis followed the protocol outlined in (30) with slight modifications. Reads were filtered per quality using Trimmomatic version 0.39 (72), trimming 11 nucleotides from the 5’ ends, utilizing a read size of 50N to minimize errors, employing a moderate penalty stiffness (0.05), trimmed at the 3’ end starting at nucleotide 89, and those shorter than 36 nucleotides were discarded. Quality control was conducted using FastQC version 0.11.8 (73). H4.V ChIP-seq were aligned to the CL Brener Esmeraldo-like strain (version 32 – TriTrypDB) using Bowtie2 version 2.3.5.1 (74) with stringent parameters (very-sensitive-local: -D 25 -R 4 -N 1 -L 19 -i S, 1,0.40 --nceil L,0,0.15). Coverage data were generated using COVERnant v.0.3.2 (https://github.com/konrad/COVERnant), resulting in ratio values derived from paired analyses of each ChIP sample versus its corresponding input. The output was saved as .wig files and visualized using IGV (http://software.broadinstitute.org/software/igv/). To identify genomic regions enriched in H4.V, coverage data were manually curated by comparing regions of enrichment in the Ty1-H4.V ChIP/input samples with those from the untagged ChIP/input control. The H4.V enriched genomic coordinates were listed in S3 Table. The ChIP-seq coverage data was analysed using a custom GFF file, annotated using a script available at (https://github.com/alexranieri/annotatePolycistron), which annotates polycistrons, dSSRs, and cSSRs. Summary plot and heatmap graphs were generated using the computeMatrix (with scale-regions and skipZeros options) and plotHeatmap (with hierarchical clustering) functions from deepTools2 (75).

### Immunofluorescence Assays (IFA) and Click-iT Reaction

For IFA, the parasites were adhered to poly-lysine-treated 8-well slides and fixed with 4% ultrapure paraformaldehyde. After fixation, slides were incubated with PBS supplemented with 20 mM glycine, permeabilized with 0.2% NP40 in PBS, blocked with 50% SFB in PBS and 3% BSA for 1 hour, then incubated with 6 M guanidine chloride in PBS, and washed. Slides were incubated for 1 hour with anti-Myc (Cell Signaling, 1:2000) and anti-Ty1 (Thermo Fisher Scientific, 1:200), followed by anti-mouse Alexa Fluor 488 (Thermo Fisher Scientific, 1:2000). Parasites expressing the Cas9 protein (Cas9 lineage) served as the control group. For S-phase detection, epimastigotes were previously incubated in complete LIT medium supplemented with 10 mM EdU for 20 minutes, as described by (36), subsequently the cells were submitted to IFA with myc or ty1 and Alexa 555 anti-mouse secondary antibodies (Thermo Fisher Scientific), and then followed for Click-iT™ EdU Imaging Kit with Alexa Fluor™ 488 (Thermo Fisher Scientific) to detect Edu-labelled DNA. All slides were mounted with Vectashield plus DAPI. Slides were viewed at 100X magnification on an inverted OLYMPUS model IX81 microscope. The cell cycle phase in IFA was determined by morphological markers based on the number of flagella, kinetoplasts, nuclei, and EdU incorporation, as described previously (36).

### Imaging Flow Cytometry (IFC)

Imaging Flow Cytometry (IFC). For assays, a total of 2x10^7^ cells in the exponential growth phase were washed with PBS and centrifuged at 1500g for 10 minutes. Subsequently, fixation was carried out using 4% paraformaldehyde (PFA) for 3 minutes, followed by washing with PBS 1x. (Note: all washes were performed by centrifugation at 1500g for 3 minutes). After fixation, cells were permeabilized with 0.1% PBS/tween-20 for 10 minutes and blocked with 3% PBS/BSA for 1 hour. Incubation with the primary antibody (anti-H3 1:2000 and anti-ty1 1:50, respectively) was conducted according to the manufacturer’s recommendations, diluted in PBS/BSA 2% for 1 hour and overnight, respectively. Following incubation with the primary antibody, three washes were performed, followed by incubation with the secondary antibody (anti-mouse PE - Texas Red 1:5000 and anti-rabbit Alexa Flour 488). Then, an incubation with the DNA marker Draq5 (1:500), diluted in PBS / BSA 2% was performed for 1h30. Finally, the cells were washed three times and stored at 4°C for subsequent analysis on the Amnis® cytometer.

### IFC data analysis

Data acquisition was conducted with 20,000 events by multispectral Image Flow Cytometry (ImageStreamx mark II Imaging flow-cytometer: Amnis Corp, Seattle, WA, Part of Luminex) using a 60× objective. The selection/gating strategy involved the exclusion of aggregates (area x aspect ratio features, spot cutting), followed by the selection of focused cells based on the Gradient RMs feature. Subsequently, three steps were applied: 1. Selection of H3-positive cells based on “Max Pixel Intensity” versus “Intensity” of H3 (Alexa Fluor 488 in channel 02); 2. From the population selected in step 1, cells in G1/S and G2/M were discriminated based on nuclear “Area Intensity” versus “Width Intensity,” combined with the kinetoplast signal (DRAQ5, channel 05). Using the imaging capabilities of ImageStream, gating was refined as follows: cells with small nuclear area, a single kinetoplast, and one flagellum were classified as G1/S; cells with enlarged nuclear area, two kinetoplasts, and two flagella were classified as G2/M.3. From cells obtained in 2, evaluation of the percentage of ^ty1-^H4.V-positive cells and their intensity levels (median) in the G1/S and G2/M populations derived from the H3-positive cells. Data were analyzed using Idea 6.3 analysis software (Amnis Corp.) and are presented as mean ± standard deviation. Student’s t test, * p <0,01.

### Salt Extraction of Chromatin-Associated Proteins

The salt extraction procedure was performed as described in (30). Approximately 10^8^ parasites (epimastigotes, metacyclics, and TCTs) expressing H2B.V^-myc^ and ^ty1-^H4.V were used and subjected to sequential salt extraction with 0, 200, 400, and 600 mM NaCl in modified RIPA buffer. The supernatant was collected and submitted to SDS-PAGE (15%) with samples normalized by DNA amount (quantified in Qubit High Sensitivity-Thermo Fisher Scientific) for subsequent Western blotting analysis.

### Western Blot (WB)

WB assays were conducted using samples from whole cell extracts (WCE) and salt gradient extractions. Membranes were blocked in a 5% solution of skimmed milk powder in TBS-T for 1 hour, followed by overnight incubation with primary antibodies at the following dilutions: anti-myc (Cell Signalling) 1:8000, anti-H3 (Abcam) 1:8000, anti-BIP 1:1000, anti-GAPDH 1:2000 (76), and anti-Ty1 (Thermo - MA5-23513) 1:500. Secondary antibodies conjugated to peroxidase (Thermo) were used at 1:3000. All washes were performed by shaking three times for 6 minutes. Membranes were visualized using ECL Western Blotting Substrate in the chemiluminescence light in Uvitec 4.7 Cambridge document photo system. Quantifications of WB results were obtained from the mean gray intensity (ImageJ). Normalizations were performed by the mean gray intensity of GAPDH labelling.

### Obtaining Telomeric Regions

The CL Brener genome was obtained through long-read sequencing, sequenced using MinION technology (Oxford Nanopore Technologies) (35). Fastq files underwent filtration using nanofilt to retain reads with a minimum quality Q score of 15 and a minimum size of 10 kb. Filtering yielded 529,882 reads with an N50 estimated at 33,447 bp. This read set was utilized to assemble a draft genome using Flye version 2.9.1-b1780 (77) with default parameters, resulting in a genome comprising 334 contigs with an average depth of 204 X, totalling 60.5 Mb and an approximate N50 of 282 kb. The genome underwent correction with two successive rounds using Medaka software (polish), culminating in the generation of a consensus genome. Telomeric regions were annotated by performing BLAST searches on contigs containing the telomeric sequence TTAGGGn (3x) of approximately 1 Kb, including the complementary reverse, against the consensus genome, yielding only one match per contig. Consequently, repeats were identified in 41 contigs. Furthermore, the Geneious Prime program (available at http://www.geneious.com/) was employed, utilizing the “similarity annotation” tool to search for TTAGGGn(3x) in the 41 detected contigs. It is noteworthy that the annotation lacks exact coordinates but represents the closest match respecting the repetition limit (TTAGGG).

## Acknowledgments

We thank Ivan Novaski Avino and Karin Navarro for technical assistance and Dr. Alex Ranieri Lima and Dr. David Pires for important input on the data and bioinformatic analysis. We thank Dr. Loyze Lima and Laboratório Estratégico de Diagnóstico Molecular from Butantan Institute for NGS analysis. We thank all TrypChrOMICs’ lab members for the many insightful discussions. This work used resources from the High-Performance Computing System of the Center for Bioinformatics and Computational Biology (NBBC) of the Butantan Institute and VeuPathDB. This manuscript was reviewed using ChatGPT for grammar, spelling, and clarity improvements. All scientific content, interpretations, and conclusions remain at the sole responsibility of the authors.

## Data availability

H4.V ChIP-seq dataset are available at Sequence Read Archive (SRA) (https://www.ncbi.nlm.nih.gov/sra) under the following accession numbers: PRJNA1124437. The genome assembly of the *T. cruzi* Cl Brener strain is deposited at PRJNA1277263.

## Supporting information

### Legends

**SF1. Identification *of T. cruzi* putative H4.V. A.** H4.V identification by phylogenic analysis. Triangle graph of the phylogenetic signal of maximum-likelihood probability calculation showing the percentages of the three topologies represented in the left triangle, referring to the tree in Fig 1A. The triangle on the right represents the attractor that, by similarity, distributes the mapped gene sequences. It was observed that the sum of the percentage of attractor vertices in the triangle on the right is greater than 60%, so the best topological model has a good phylogenetic signal, which is why it was used in the generated phylogenetic tree (ultrafast bootstrap values by 1000 replicates - best-fit model: TN+F+G4). **B.** Canonical histone H4 and putative histone H4.V from *T. cruzi* (CL Brener strain – S-and P-haplotype). Nucleotide and amino acid alignments were performed at Clustal Omega Clustal Omega - Multiple Sequence Alignment from EMBL-EBI plotted on Geneious Prime software. Nucleotides and amino acids highlighted in colour indicate mismatches in the alignment.

**SF2. ^ty1-^H4.V parasites generated by CRISPR-Cas9. A.** Scheme representing the H4.V locus before (top) and after (bottom) gene editing by CRISPR. The donor DNA has three copies of ty-1 sequences; the 5’ UTR of *T. brucei* actin, 3’UTR of aldolase and the 5’ UTR of aldolase (dark grey); the blasticidin resistance gene (red); and homology arm (Harm-dark blue) that are homologous to the 5’ UTR of H4.V. The red arrows indicate the primers that were used to validate the genome edition, and black arrows indicate the gene transcriptional direction. **B**. PCRs of ^ty1-^H4.V parasites using the indicate pair of primers. Amplicons were fractionated by 1% agarose gel showing the insertion of the ^ty1-^tagged specifically at the H4.V locus. The blasticidin gene and ty-1 were detected, ∼1185 bp (primers 1+4) and ty-1 (primers 2+4 e 2+3), respectively. Primer 4 was designed to anneal specifically to H4.V regions, with no complementarity to canonical H4. The genomic DNA of parasites expressing Cas9 were used as negative control (C-). **C.** Western blot assay using anti-ty1 confirmed the H4.V expression in epimastigotes clones. Non-transfected parasites were used as negative control (Ctl). **D.** Parasites expressing Cas9 and ^ty1-^H4.V growth curves (log10). Unpaired T-test of area under the curve, p = ns. **E.** Metacyclic trypomastigotes expressing Cas9 or ^Ty1-^H4.V were quantified in RPMI culture supernatants after 8 days. Unpaired T-test, p = ns (p=0.1609). The experiments were performed in biological triplicates. ns: no statistically significant differences.

**SF3. Distribution and validation of H4.V enrichment.** Genomic features flanking cSSRs without (A) or with (B) H4.V enrichment. cSSRs located “between mDNAs and tDNAs” – p-value< 0.00115 (Chi-square test with simulation of 1,000 replications) with adjusted p-value = 0.00625. **C.** Plot Enrichment graph (from deepTools) from all H4.V manually enriched genomic regions confirms higher enrichment of H4.V in all ^Ty1-^H4.V ChIP sample replicates compared to control samples (input control-Cas9, ChIP control-Cas9, input ^Ty1-^H4.V). **D.** Quantification between enriched genomic coordinates from ChIP and input samples described in (C.). Multiple comparisons One-way ANOVA were performed in which * (p<0.05) and ** (p < 0.005).

**SF4. H4.V enrichment in chromosome ends. A**. IGV screenshot of ^Ty1-^H4.V ChIP-seq data from the chromosome 23 end. The track “ty1-H4.V-epi”, “untagged control”, “H2B.V-myc” and “H3” represent *.wig files from the ratio values (ChIP/input) obtained by COVERnant (window of 1001 bases per step of 201). The bed tracks in green represent the conserved genome compartment, in red the disruptive compartment, and in yellow genes RHS. Dashed box represent enrichment of histone H4.V at chromosome ends. **B.** Heatmaps of ChIP-seq data from two replicates ^Ty1-^H4.V (ChIP/input) and untagged control parasites in 51 telomeric regions from 41 contigs. The telomeric coordinates from the CL Brener genome assembly obtained from long reads. The matrix was created with the scale-regions option that fits all regions of interest (telomeres) within Start (S) and End (E). Regions 10 kb downstream and upstream of telomers were also evaluated. Note that, due to the characteristic location of telomeres at chromosome ends, no H4.V enrichment (zero – dark blue) is expected either upstream or downstream of telomeric regions across all 41 contigs. In contrast, the telomeric regions themselves (between S and E) display higher levels of H4.V enrichment (red).

**SF5. IFC analysis of ^ty1-^H4.V parasites. A.** Fluorescence intensity plots showing the following labels: Alexa Fluor 488 (histone H3) in channel 02 (ch02); DRAQ5 (nucleus) in channel 05 (ch05), PE-Texas Red (histone H4.V) in channel 04 (ch04). **B.** Sequential gating strategy to extract H4.V abundance on different cell cycle phases. Initial gate selection of histone H3-positive cells based on “Max Pixel Intensity” versus “Intensity of H3” (top left); followed by cell cycle phase discrimination (G1/S and G2/M) based on nuclear Area Intensity versus Width Intensity (DRAQ5 detection) combined with kinetoplast/flagella quantification (top right); subsequent selection of ^ty1-^H4.V-positive cells (“Max Pixel Intensity” versus “Intensity”) within the H3-positive populations in each cell cycle phase (bottom quadrants). **C.** Selected images of triple-stained epimastigotes labelled for histone H3, histone ^ty1-^H4.V, and nuclear staining, with corresponding merged images across cell cycle phases obtained from IFC. **D.** Graph illustrating the percentage of positive cells for both H3 and H4.V in each cell cycle phase. Graphs and images were generated using the Idea 6.3 analysis software (Amnis Corp.).

